# Menaquinone-specific oxidation by *M. tuberculosis* cytochrome *bd* is redox regulated by the Q-loop disulfide bond

**DOI:** 10.1101/2024.01.09.574813

**Authors:** Tijn T. van der Velden, Caspar Y. J. Waterham, Lars J. C. Jeuken

## Abstract

Cytochrome *bd* from *Mycobacterium tuberculosis* (*Mtbd*) is a menaquinol oxidase that has gained considerable interest as an antibiotic target due to its importance in survival under infectious conditions. *Mtbd* contains a characteristic disulfide bond that has been hypothesized to confer a redox regulatory role during infection by constraining the movement of the menaquinone-binding Q-loop. Interference of reductants used in the standard activity assay of quinol oxidases has prevented testing of this hypothesis. Here, the role of the disulfide bond and quinone specificity of *Mtbd* has been determined by the reconstitution of a minimal respiratory chain consisting of a NADH dehydrogenase and *Mtbd*, both in detergent and native-like lipid environments. Comparison to cytochrome *bd* from *Escherichia coli* (*Ecbd*) confirms that *Mtbd* is under tight redox regulation and is selective for menaquinol, unable to oxidize either ubiquinol or demethylmenaquinol. Reduction of the *Mtbd* disulfide bond resulted in a decrease in oxidase activity up to 90%, depending on menaquinol concentrations. In addition, the catalytic rates of *Ecbd* and *Mtbd* are over 10 times lower with the natural lipophilic quinones in comparison to their often-used hydrophilic analogs. Additionally, unlike *Ecbd*, the activity of *Mtbd* is substrate inhibited at physiologically relevant menaquinol concentrations. We signify *Mtbd* as the first redox sensory terminal oxidase and propose that this enables *Mtbd* to adapt its activity in defence against reactive oxygen species encountered during infection by *M. tuberculosis*.

## Introduction

Cytochrome *bd* (cyt *bd*) is a terminal oxidase found in the respiratory chain across various bacterial phyla and couples the oxidation of quinols to the reduction of molecular oxygen to water^1–3^. The chemical protons released by quinol oxidation are separated from the proton uptake for oxygen reduction by the periplasmic membrane, generating a proton motive force required for ATP synthesis^3^. Compared to other terminal oxidases, cyt *bd* is distinguished by its resistance to inhibitors such as cyanide, high affinity for oxygen and upregulation under microaerobic conditions.

Recent findings highlight the significance of cyt *bd* as a key survival factor for *Mycobacterium tuberculosis* during infection, particularly in the hostile granulomas where the bacteria reside. When *M. tuberculosis* cyt *bd* (*Mtbd*) functions as the sole terminal oxidase, it can maintain a bacteriostatic state^4,5^, and enhances resistance against numerous antibiotics, reactive oxygen species, and other toxic compounds^6–11^. This pivotal role during infection has identified *Mtbd* as a potential antibiotic target.

Despite this therapeutic potential, most of our current understanding of *Mtbd* at the molecular level is obtained from studies performed on homologous enzymes, such as *Corynebacterium glutamicum* cyt *bd* and the two *Escherichia coli* cyt *bd* isoforms, cyt *bd*-I (*Ecbd*) and cyt *bd*-II. Since these enzymes have been shown to have distinct structural features, number of subunits, and substrate binding pockets^12–16^, caution is required when associating prior knowledge to *Mtbd*. An important characteristic feature of *Mtbd* is a disulfide bond constraining the n-terminal Q-loop domain near the quinone binding pocket. It has been hypothesized that this disulfide bond acts as a gatekeeper for the canonical quinone binding pocket close to heme b_558_, and might confer a regulatory function^12,17^. Redox-sensing disulfide bonds have been found in other proteins such as transcription factors, signalling enzymes, CO_2_ reductases, and peroxidases as a defence against oxidative stress^18,19^. However, whether such a regulatory function is conferred by the *Mtbd* Q-loop remains speculative, because its validation has been hampered by the need for chemical reductants to reduce the quinone substrates in the standard activity assays for cyt *bd* oxidase. Given its significance as a potential antibiotic target, it is essential to uncover the main features of *Mtbd* enzyme kinetics.

*Ecbd, Mtbd* and cyt *bd*’s from other species use distinct quinone subtypes for turnover. The main bacterial quinone subtypes include menaquinone (MK), demethylmenaquinone (DMK), and ubiquinone (UQ), each differing in their quinone headgroup structure and corresponding redox potential (Fig 1A)^20^. In addition, these quinones contain a hydrophobic isoprenoid tail made from a distinct number of isoprene units^21^. By convention, the number of isoprene units is indicated by the number (n) after the quinone, e.g. MK-n. *M. tuberculosis* primarily relies on MK-9^22^, while *E. coli* demonstrates distinct functionalities for MK-8, DMK-8, and UQ-8^23,24^. Moreover, *E. coli* upregulates MK-8 during microaerobic conditions, coinciding with the upregulation of *Ecbd*, highlighting the adaptability of the *E. coli* quinone pool composition based on environmental requirements^20,25^.

**Figure 1.**
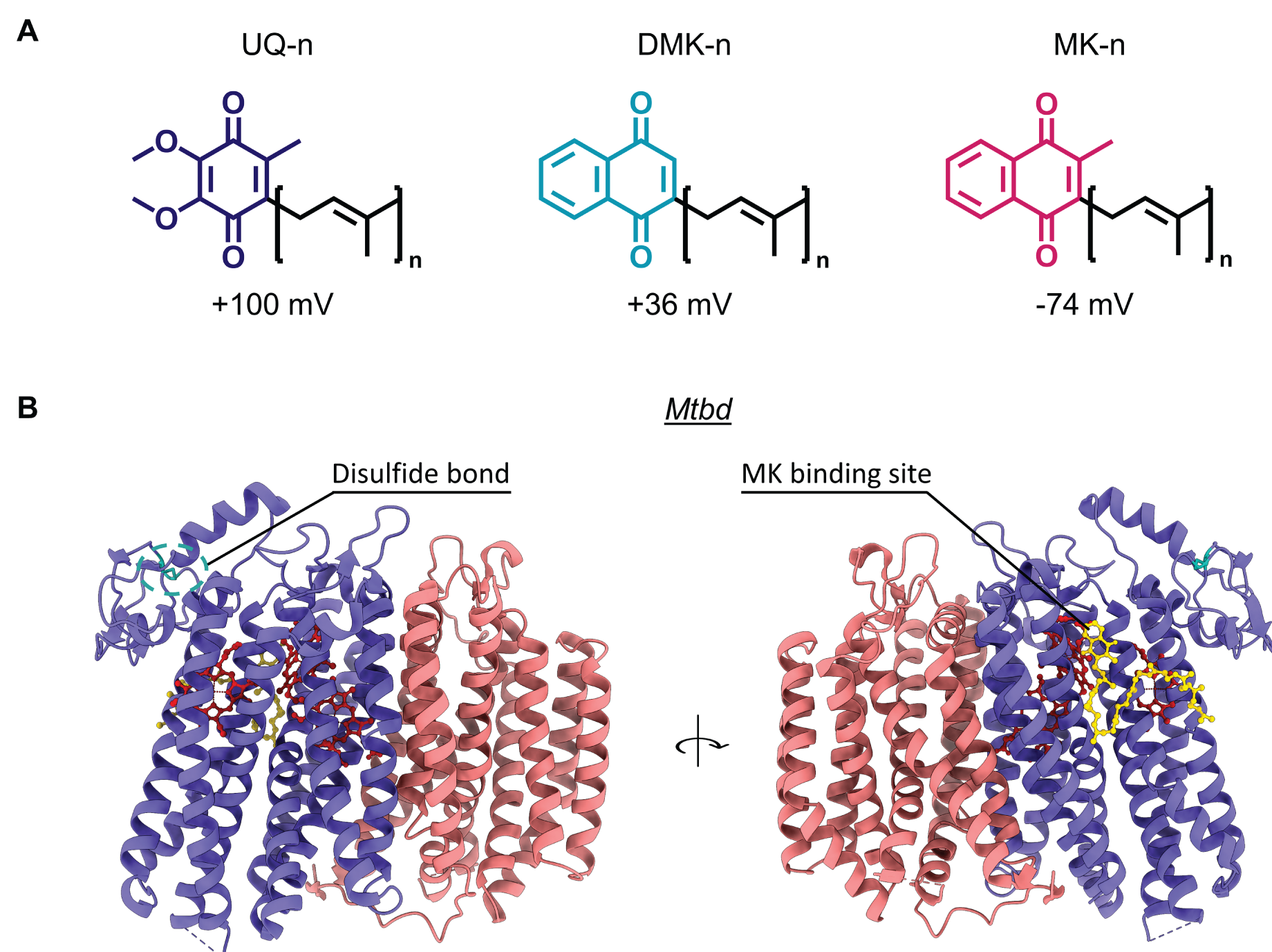
Overview of quinone subtypes and structural features of *Mtbd*. (**A**) The structure of quinone analogs used in this study with the number of isoprenoid units (n) and their reduction potentials^20^. (**B**) Structure of *Mtbd*, CydA (purple), CydB (orange), (PDB: 7NKZ) indicating the Q-loop disulfide bond and MK binding site.

The interactions between quinones and cyt *bd* are intricate, as demonstrated by the distinct quinone binding sites in *Ecbd* and *Mtbd* (SI Fig 1). While *Ecbd* has a proposed quinone binding pocket at the Q-loop of subunit CydA, transferring electrons to closely located heme *b*_558_^14^, *Mtbd* shows quinone binding at the back of CydA, which transfers electrons directly to heme *b*_595_, bypassing the initial heme *b*_558_ (Fig 1B)^12^. Additional quinone binding pockets are found that are spatially separated from all three hemes, prohibiting participation in quinol oxidation. For instance, *Ecbd* contains UQ-8 in its CydB subunit, thought to confer structural stability^14^ or allosteric regulation, where exchanging different quinone subtypes may alter enzyme

Neither hypothesis has yet been unambiguously proven. Although for *E. coli* cyt *bd*-II and *C. glutamicum* cyt *bd*, preincubation with MK has been shown to increase enzymatic turnover^27^, single-particle cryo-EM studies were unable to indicate that MK incubation resulted in an exchange of this ‘structural’ quinone’^16^. Hence, how the isoprenoid tail and redox properties of these quinone subtypes affect the kinetics of *Ecbd* and *Mtbd* requires further study.

Most studies make use of detergent solubilized cyt *bd* with water soluble quinone analogs. The use of detergent micelles, although convenient for studying cyt *bd* in solution, introduces an artificial environment that deviates from the native conditions. These detergent micelles can alter the conformation, partially unfold proteins, and strip away vital bound or annular lipids, leading to a loss of native mechanistic function and kinetic rates^28^. Additionally, using water soluble quinone analogs could significantly alter enzyme kinetics, as indicated when the quinone isoprenoid tail layout was shown to critically affect respiratory efficiency in *M. tuberculosis in vivo*^29^. The thorough evaluation of cyt *bd* quinone interactions and disulfide regulation requires a native membrane environment and natural quinone substrates to mimic *in vivo* conditions.

To study the specific interaction of *Ecbd* and *Mtbd* with the different quinones and interrogate the role of the *Mtbd* Q-loop disulfide bond, a minimal respiratory chain was constructed in which cyt *bd* is combined with a NADH-quinone oxidoreductase, *Caldalkalibacillus thermarum* NDH-2 (Figure 2). This revealed that water soluble quinone analogs lead to vast overestimations of cyt *bd* turnover rates, highlighting the need for precise experimental design. Furthermore, In contrast to *Ecbd, Mtbd* is subject to substrate inhibition by MK-9 at physiologically relevant concentrations.

**Figure 2.**
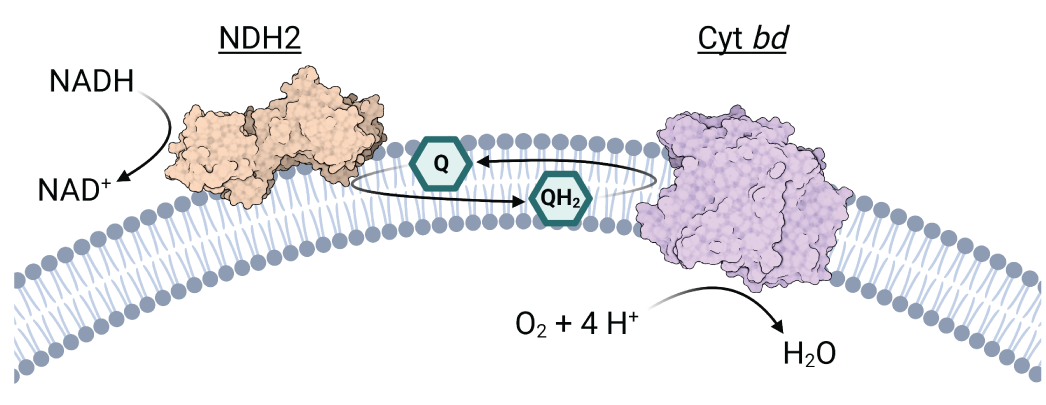
A minimal respiratory chain proteoliposomal system to interrogate specific interactions between quinones and cyt *bd*. This proteoliposomal system can also be used in detergent.

Additionally, we signify *Mtbd* as the first terminal oxidase with a redox sensory disulfide bond. We propose that this enables *Mtbd* to adapt its activity to environmental redox pressures in defence against reactive oxygen species encountered by *M. tuberculosis* during infection.

## Results

### Cyt *bd* oxygen consumption kinetics with water soluble quinone analogs

*Ecbd, Mtbd* and NDH-2 were expressed and purified via affinity and size exclusion chromatography (SI Fig 2). A minimal respiratory chain was constructed with an excess of NADH and NDH-2, such that the quinone pool remains fully reduced. Enzyme activity of *Ecbd* and *Mtbd*, comparing different quinone substrates, was determined by monitoring oxygen consumption in a Clark electrode setup. Previous reports indicate that quinols, especially menaquinols, auto-oxidize under aerobic conditions in a concentration dependent manner^30^. To account and correct for the quinol auto-oxidation rate, the oxygen consumption was quantified for each assay before the addition of cyt *bd* (Fig 3A). Activity was first measured in detergent conditions using water-soluble quinone analogs. *Ecbd* displayed standard Michaelis-Menten kinetics with a 6-fold lower K_m_ for MK-1 compared to UQ-1 (Fig 3B, Table 1). K_cat_ values for the three quinone analogues were similar (Table 1) ranging from 736 to 1033 Q/s (Table 1).

**Table 1.** Kinetic parameters for the oxidation of UQ-1, DMK-1, MK-1 by *Ecbd* ^a^.

| $K_{cat}$ (Q s<sup>-1</sup>)<sup>b</sup> | | | $K_m$ (μM)<sup>c</sup> | | | $K_{cat}/K_m$ (x10<sup>-6</sup> M<sup>-1</sup> s<sup>-1</sup>)<sup>d</sup> | | |
| --- | --- | --- | --- | --- | --- | --- | --- | --- |
| UQ-1 | DMK-1 | MK-1 | UQ-1 | DMK-1 | MK-1 | UQ-1 | DMK-1 | MK-1 |
| 1033 ± 85 | 736 ± 51 | 846 ± 38 | 149 ± 26 | 44 ± 10 | 22 ± 4 | 6.9 ± 1.3 | 16.7 ± 4.0 | 38.5 ± 7.2 |

<sup>a</sup> $K_{cat}$ values are represented in quinones per second (2 e<sup>-</sup>/Q). Oxygen consumption measurements were performed in 50 mM MOPS, 150 mM NaCl, 0.005% LMNG, pH 7.0 (n=3). Data represents best Michaelis-Menten fit values with the SEM. <sup>b</sup> UQ-1 vs DMK-1 \*. <sup>c</sup> UQ-1 vs DMK \*\*, UQ-1 vs MK-1 \*\*\*, DMK-1 vs MK-1 \*. <sup>d</sup> Errors represent the propagated SEM. \* p< 0.05, \*\* p< 0.001, \*\*\* p<0.0001

**Figure 3.**
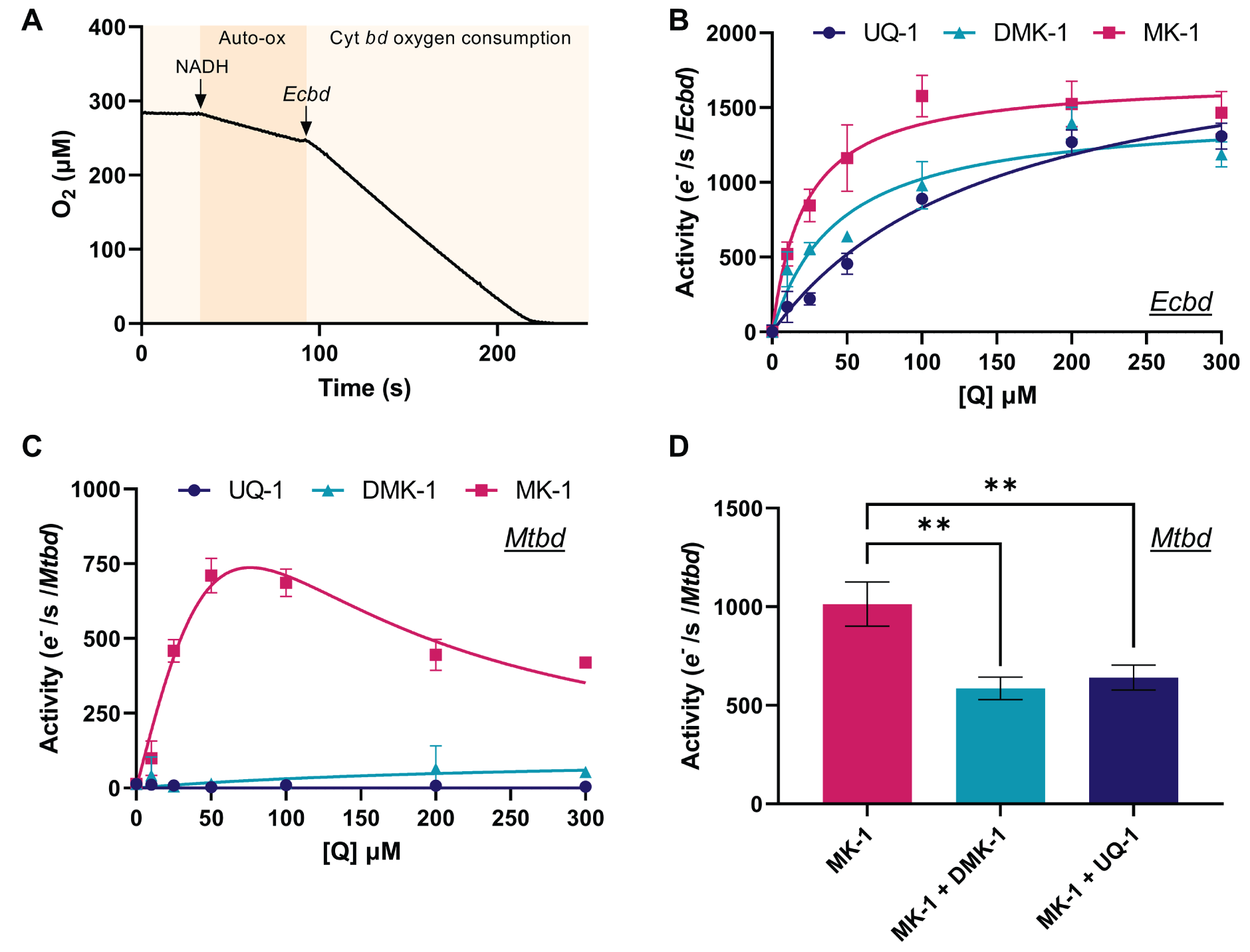
Oxygen consumption kinetics of detergent solubilized cyt *bd* with different quinone substrates. (**A**) Example of an oxygen consumption trace with MK-1 (50 μM) auto-oxidation (Auto-ox) and *Ecbd* oxygen consumption indicated. (**B**) *Ecbd* and (**C**) *Mtbd* oxygen consumption kinetics using the assay described in the text. Lines represent Michaelis-Menten fits without (*Ecbd*, see Table 1 for parameter values) or with (*Mtbd*) substrate inhibition. (**D**) *Mtbd* activity with 50 μM MK-1 in the absence or presence of either DMK-1 or UQ-1 (both 300 μM). Data is represented as the average and standard deviation of triplicate measurements of two different protein preparations (n=3). ** p<0.01

The kinetic profile of *Mtbd* was profoundly different than that of *Ecbd* (Fig 3C). *Mtbd* exclusively turns over MK-1 and is subject to substrate inhibition. While substrate inhibition has been observed in other oxidases^16,31,32^, its molecular mechanism remains undefined. Multivariate fits of the observed substrate inhibition profile indicated a large interdependency of parameters (K_m_; V_max_; inhibition constant, K_i_) preventing the determination of the kinetic parameters for MK-1 oxidation in spite of the good fit (Fig 2C). Importantly, *Mtbd* failed to initiate turnover with either DMK-1 or UQ-1. To investigate whether this inactivity resulted from a lack of substrate binding or a thermodynamic barrier limiting quinol oxidation, the binding of UQ-1 and DMK-1 to *Mtbd* was assessed. It was reasoned that DMK-1 and UQ-1 will present as competitive inhibitors if they bind to the same active-site pocket as MK-1. Upon addition of an excess DMK-1 or UQ-1, oxygen consumption activity was indeed significantly inhibited, suggesting that DMK-1 and UQ-1 bind to the active site, but are unable to be oxidized by *Mtbd* (Fig 3D). To determine whether the observed substrate specificities in detergent are indicative of *in vivo* conditions, the same principles were applied to a proteoliposomal system where cyt *bd* can be studied in a native lipid environment using the long isoprenoid quinones MK-9 and UQ-10.

### *Ecbd* and *Mtbd* oxygen consumption in proteoliposomes

*Ecbd* and *Mtbd* were reconstituted in a proteoliposomal system that emulates a minimal respiratory chain using the same enzymatic components used in our detergent setup. In this system, oxygen consumption is used to monitor cyt *bd* activity, while excess NDH-2 binds the proteoliposomes to facilitate the reduction of the membrane-embedded quinone pool upon the addition of NADH (Fig 2).

*Mtbd* and *Ecbd* were reconstituted in liposomes containing 0.25% - 1% (w/w; quinone/lipid) of the desired quinone (MK-9 or UQ-10) to achieve maximum catalytic rates. Quinone concentrations in bacterial membranes are estimated between 0.3 – 1.5% (w/w) (see SI). Furthermore, 1% MK-9 or UQ-10 approximately coincides with a membrane concentration in the order of 10-15 mM (see SI), well above the K_m_ determined for *Ecbd*. We note however, that for the water soluble quinone analogs an unknown equilibrium exists between quinones in solution and quinones in detergent micelles, making the two systems difficult to directly compare. Consistent with the UQ-1/MK-1 kinetics, *Ecbd* showed a marginally higher catalytic rate with UQ-10 than MK-9 (Fig 4A). Additionally, *Mtbd* displayed the same specificity for MK turnover, confirming the quinone analog data (Fig 4B). Substrate inhibition was confirmed by increasing the MK concentration from 0.25% to 1%, resulting in a significant decrease in enzyme activity.

**Figure 4.**
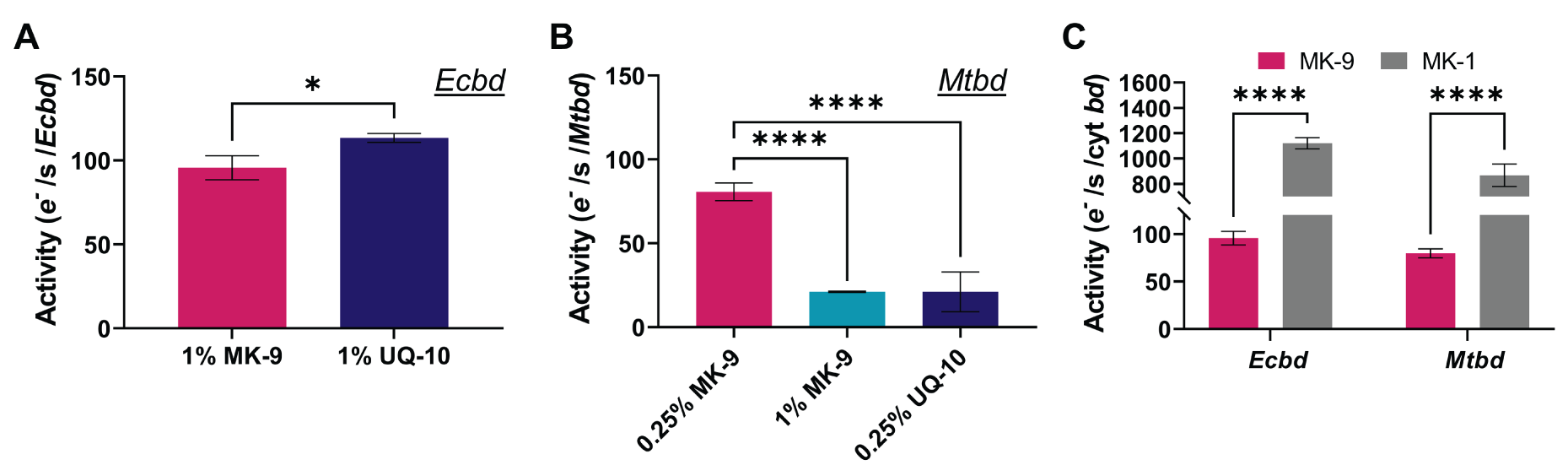
Quinone substrate specificity of *Ecbd* and *Mtbd* in a native lipid environment. (**A**) Catalytic rates of *Ecbd* (20 nM) proteoliposomes containing 1% (w/w) of either UQ-10 or MK-9. (**B**) Catalytic rates of *Mtbd* (20 nM) with different concentrations of either UQ-10 or MK-9. (**C**) Reconstituted *Ecbd* and *Mtbd* activity with native MK-9 (*Ecbd* 1% (w/w) MK-9, *Mtbd* 0.25% (w/w) MK-9) versus MK-1 analog (50 μM). Data is represented as the average and standard deviation from three individual proteoliposome reconstitutions. * p<0.05, **** p<0.0001

To study the effect of the quinone isoprenoid chain on cyt *bd* activity, *Ecbd* and *Mtbd* were reconstituted in proteoliposomes and activated with either membrane-embedded MK-9 or water soluble MK-1. Surprisingly, the isoprenoid chain length of the quinone was shown to have a significant effect on the maximum catalytic rate of cyt *bd* (Fig 4C). Utilizing the natural membranous quinones resulted in tenfold lower catalytic rates, emphasizing the role of the isoprenoid tail in substrate binding, via either diffusion or binding kinetics. This highlights that previously reported catalytic rates using quinone analogs substantially overestimate the catalytic rates achieved *in vivo*.

### *Mtbd* activity is regulated by a unique Q-loop disulfide bond

*Mtbd* is characterized by a unique disulfide bond that was hypothesized to reduce the flexibility of the substrate binding Q-loop (Fig 5A) and thereby confer a regulatory role for enzyme activity^12,33^. Previous studies faced limitations examining this mechanism, as they relied on chemical reductants such as DTT to reduce the quinone pool, which inadvertently affects the disulfide bond itself. Using NDH-2 to enzymatically reduce the quinone pool enables the comparison of *Mtbd* activity before and after incubation with chemical reductants to reduce the disulfide bond and probe its effects on *Mtbd* activity.

**Figure 5.**
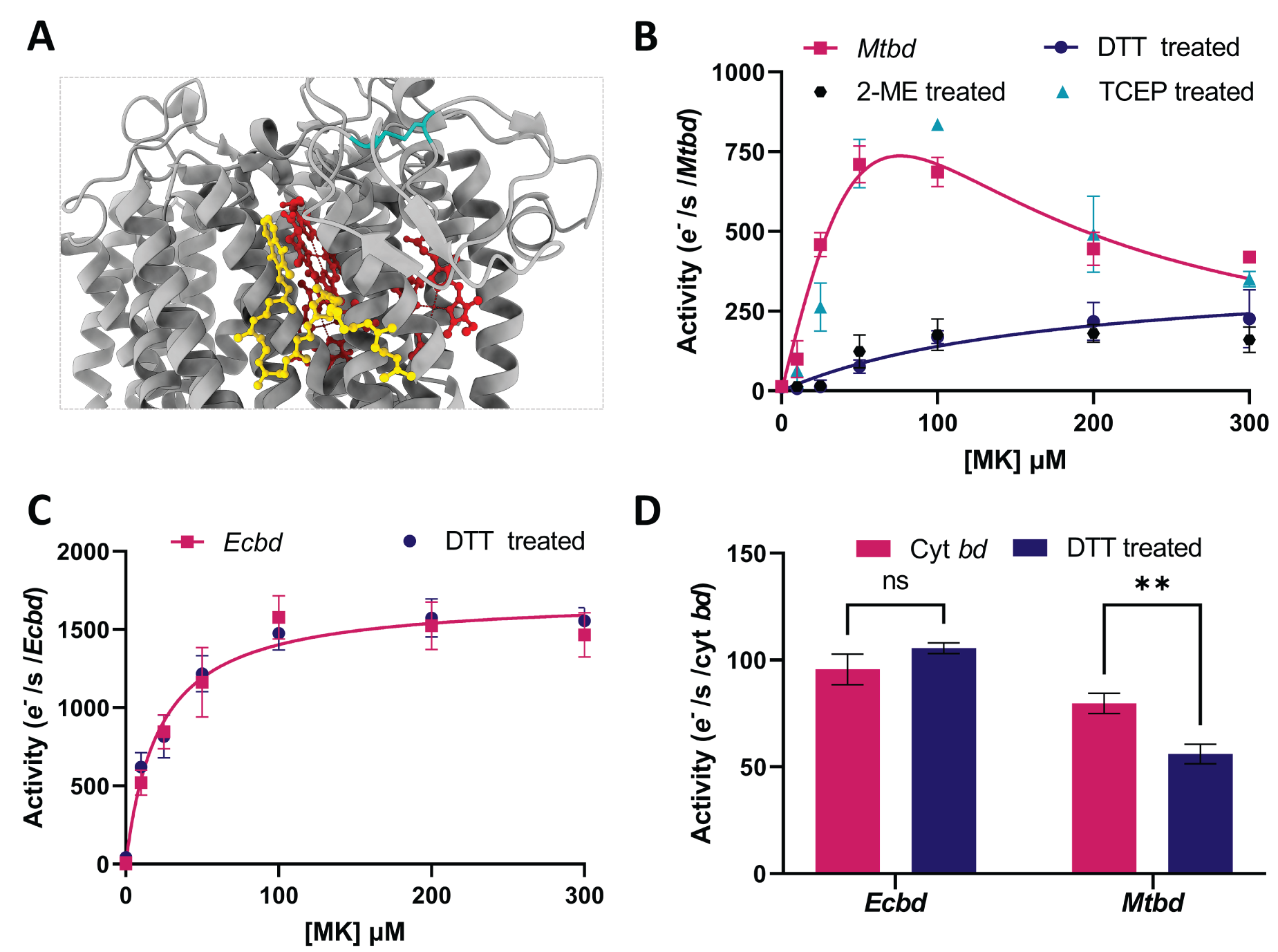
*Mtbd* regulation by the Q-loop disulfide bond. (**A**) Cryo-EM structure of *Mtbd* (PDB: 7NKZ) with heme groups (red) MK-9 (yellow) and unique Q-loop disulfide bond (blue). The structure was analyzed using ChimeraX^34^ (**B**) *Mtbd* MK-1 kinetics in detergent micelles before and after treatment with TCEP, DTT, or 2-ME. Significant inhibition of oxidase activity is observed after DTT or 2-ME treatment. (**C**) *Ecbd* MK-1 kinetics in detergent micelles before and after DTT treatment. No difference is observed. (**D**) Oxidase activity of proteoliposomes containing *Mtbd* (0.25% MK-9) or *Ecbd* (1% MK-9) before and after DTT treatment. ** p<0.01

Incubation of *Mtbd* with either DTT or 2-mercaptoethanol (2-ME) showed a significant reduction in oxidase activity (Fig 5B). In addition to the decrease in *Mtbd* activity, the disulphide bond breakage eliminates MK substrate inhibition, strongly suggesting the reduction of the disulfide bond changes the structure or dynamics of the quinone binding pocket. Unexpectedly, preincubation with the reductant tris(2-carboxyethyl)phosphine (TCEP) did not reduce *Mtbd* activity, indicating that interactions with the disulfide bond might be hindered by surface accessibility or charge repulsion (SI fig 3). To verify that the chemical reductants only impact the disulfide bond and no other aspects of *Mtbd*, the same measurements were performed with *Ecbd*, which lacks the Q-loop disulfide bond (Fig 5C). This confirms that there is no difference in *Ecbd* activity after treatment with DTT and consolidates the conclusion that the decrease in the activity of *Mtbd* stems from the reduction of the Q-loop disulfide bond. To ensure that the observed effects are not artifacts of the detergent environment, the measurements were repeated in our proteoliposomal system (Fig 5D). Again, the *Ecbd* activity remains unchanged by the DTT treatment, while the *Mtbd* activity significantly decreases. The DTT-mediated decrease in proteoliposomal *Mtbd* is less pronounced in comparison to the detergent solubilized *Mtbd*. We ascribe this to the fact that the proteoliposome system has higher quinone concentrations, for which the difference is less pronounced (Fig 5B).

## Discussion

Cyt *bd* is a critical part of the prokaryotic respiratory chain to maintain ATP regeneration under microaerobic conditions^3^. *Mtbd* has been highlighted for its essentiality under oxygen limiting conditions, and its related interest as an antibiotic target against *M. tuberculosis*. Most current knowledge, however, comes from mechanistic studies on homologous enzymes such as *Ecbd* and *C. glutamicum* cyt *bd*, which have diverse structural features, such as a lack of the characteristic *Mtbd* Q-loop disulfide bond ^12–15^.

The minimal respiratory chain, both in detergent and liposomes, unambiguously indicates that *Ecbd* can oxidize UQ, MK and DMK. The affinity of *Ecbd* for UQ-1 (k_m_: 149 ± 26 μM) is in line with previously reported values^31,32,35–37^, while the catalytic rate at 200 μM UQ-1 (1268 ± 81 *e*^*-*^ s^-1^), is slightly higher than literature reported values (889 ± 30 *e*^*-*^ s^-1^)^14^ and is attributed to the difference in the detergent environment and assay conditions. The fact we see activity with MK is in direct contrast with an earlier study that suggests that *Ecbd* cannot oxidize MK^16^. We note that in this earlier study, MK was reduced by DTT. Indeed, when we repeated the assay with DTT instead of NADH/NDH-2 to reduce MK, we did not observe any activity. Here, we propose that although DTT is a good reductant for UQ and hence a good reductant to measure oxygen-reducing activity by cyt *bd*, DTT might be a poor reductant and rate-limiting when the assay is performed with MK. The observation that *Ecbd* can oxidize MK and DMK is consistent with the *in vivo* upregulation of MK and DMK during microaerobic respiration, conditions that also lead to increased expression of *Ecbd*^24,25,38^.

The catalytic rates of both *Ecbd* and *Mtbd* were approximately ten-fold lower when using the natural quinones in comparison to their often-used water-soluble analogs. Similar effects have been shown before, where small differences in quinone isoprenoid units altered enzyme activity^27,39^. This highlights the need for physiologically relevant assay conditions to obtain appropriate kinetic rates and inhibition values. The difference in activity when using the native quinones could be explained by a difference in binding or diffusion kinetics, as many of the quinone-bound oxidase structures have shown that the quinone isoprenoid tails play a significant role in substrate-enzyme interactions^12,40^, and longer chains might result in slower substrate binding and release^41^.

*Mtbd* exclusively catalyzes MK oxidation, while competition experiments suggest that UQ and DMK can bind to the same active site. A similar phenomenon was shown for cyt *bd* from *C. glutamicum*, and cytochrome *bcc* from *M. tuberculosis*, which also lacked oxidase activity using UQ as a substrate ^27,30^. We postulate that this lack of UQ-1 turnover in *C. glutamicum* cyt *bd* is caused by a thermodynamic barrier between two-electron oxidation of UQ (100 mV) and the single-electron acceptor heme *b* (102 mV) ^13,42^. Similar suggestions have been made for mycobacterial cytochrome *bcc*, where the heme redox potentials are too low to allow the oxidation of UQ^30,43^. Although the heme potentials for *Mtbd* are unknown, we postulate that the same hypothesis applies here.

*Mtbd* was unable to oxidize DMK, which is similar in structure to MK and has a reduction potential that lies in between that of MK and UQ. In *M. tuberculosis*, DMK is a precursor in MK biosynthesis. Interestingly, small molecule inhibitors of MenG, the enzyme that converts DMK into MK, were bactericidal in *M. tuberculosis*^33,44^. This substantiates the inability of *Mtbd* to turn over DMK and highlights the tight redox control between the quinone pool and *Mtbd*. Additionally, this emphasizes inhibition MK biosynthesis as a valid antibiotic strategy that targets the fundamental electron transfer steps between the respiratory chain components.

In contrast to our observation reported here, *Mtbd* expressed in *E. coli* has been reported to oxidize UQ when assayed in crude membrane extracts^39^. Potentially, the observed UQ oxidation was caused by interfering oxidases present in the membrane extracts, such as *E. coli* cyt *bd*-II. Further studies would be required to determine why the assay in crude *E. coli* membrane extracts gives rise to different results than enzymes purified from *M. smegmatis*, as reported here.

Importantly, the *Mtbd* disulfide bond is unique among the currently studied cyt *bd* oxidases. Despite the conservation of these cysteine residues in *M. smegmatis*^45^ and *C. glutamicum*^13^ cyt *bd*, these proteins do not show a formed cysteine bond in their structures, suggesting that the bond is not necessary for the protein to remain in a folded state. Molecular dynamics studies have indicated that the *Mtbd* disulfide bond can alter Q-loop flexibility, potentially regulating *Mtbd* activity^12^. Regretfully, none of the aforementioned proteins have structures solved in both the formed and broken disulfide bond states. Here we show that the chemical reduction of the *Mtbd* disulfide bond results in a significant decrease in oxidase activity. To our knowledge, this is the first proof of a terminal oxidase under the regulation of a redox sensing disulfide bond. Remarkably, when *M. smegmatis* cyt *bd* was studied and found to have a broken disulfide bond, it also presented low catalytic rates (21.6 ± 2.8 e^-^ s^-1^)^45^, significantly lower than the catalytic rates we report here. Potentially, the *M. smegmatis* disulfide bond is reduced during isolation or only gets formed under the oxidative pressure of reactive oxygen species. It would be interesting to assess if the activity of *M. smegmatis* and *C. glutamicum* cyt *bd* increases after the formation of the disulfide bond, for instance by incubation with copper or reactive oxygen species such as hydrogen peroxide.

Interactions of *Mtbd* with environmental redox pressures might be important as *M. tuberculosis* can survive in reactive oxygen species rich environments during infection^46^, and *Mtbd* has a suggested role in hydrogen peroxide resistance^47^. In addition, *Ecbd* has been shown to actively detoxify the cellular environment from hydrogen peroxide^48^ and peroxynitrite^49^, indicating potential additional roles for *Mtbd* as a survival factor for *M. tuberculosis*. Future research should focus on the physiological role of the *Mtbd* disulfide bond and its influence on *M. tuberculosis* survival under various conditions. Additional investigations of the interplay between *Mtbd* and environmental redox pressures, especially regarding its potential detoxifying role as seen with *Ecbd*, will offer valuable insights into the adaptive role of *Mtbd* in the defence against reactive oxygen species and antibiotic compounds encountered during infection.

## Experimental procedures

### Expression and purification of *E. coli* cytochrome *bd*-I

The expression of *E. coli* cytochrome *bd*-I (*Ecbd*)was performed as previously described ^50^, with slight modifications. Briefly, MB43 cells ^51,52^ transformed with pET17b-CydABX-linkerstreptag were grown overnight in LB with 100 μg/mL ampicillin (250 RPM, 37°C). The culture was diluted to OD ∼0.1 in LB ampicillin and grown to OD∼ 0.4. Expression was induced by the addition of 0.45 mM IPTG and carried out until OD ∼2.0. Cells were harvested by centrifugation (6371 rcf, 20 min, 4°C) and resuspended in 50 mM MOPS, pH 7.4, 100 mM NaCl, cOmplete™ EDTA-free Protease Inhibitor (ROCHE), at 1 g wet cells per 5 mL buffer. Cells were disrupted by a single pass through a Stansted pressure cell homogeniser (270 MPa). Unbroken cells were pelleted and discarded by centrifugation (10.000 rcf, 20 min, 4°C). Crude membranes were isolated by ultra-centrifugation (200.000 rcf, 1h, 4°C) and resuspended to 10 mg/mL protein concentration in 50 mM MOPS, 100 mM NaCl, pH 7.4. Detergent extraction of the membrane proteins was performed by incubation with 0.5% Lauryl maltose neopentyl glycol (LMNG) for 1 hour at 4°C with gentle mixing. Insoluble material was pelleted and discarded by ultra-centrifugation (200.000 rcf, 30 min, 4°C) followed by application of the soluble fraction to a StrepTrap HP column (Cytiva) at 1 mL/min. To remove unbound proteins, the column was washed with 50 mM sodium phosphate, 300 mM NaCl, 0.005% LMNG, pH 8.0. Elution was performed by the addition of 50 mM sodium phosphate, 300 mM NaCl, 2.5 mM desthiobiotin, 0.005% LMNG, pH 8.0, after which purity was confirmed by SDS-page. Fractions containing pure *E. coli* cytochrome bd-I were pooled, concentrated, and stored at -80°C until further use.

### Expression and purification *M. tuberculosis* cytochrome *bd*

*M. tuberculosis* cytochrome *bd* (*Mtbd*) expression and purification were performed as previously described with slight modifications ^53^. *M. smegmatis* MC^2^ 155 ΔCydAB ^54^ was transformed with the pLHCyd plasmid using electroporation with an Eppendorf eporator (2.5 kV). Positive transformants were selected by plating the cells on LB agar with 50 μg/mL hygromycin. A starter culture was inoculated in LB-hygromycin and grown for 72 hours (250 RPM, 37°C). The culture was diluted 1:100 and grown for an additional 72 hours (200 RPM, 37°C) until the cells were harvested (6371 rcf, 20 min, 4°C) and resuspended in a 5-fold volume of 50 mM Tris-HCl (pH7.4), 5 mM MgCl_2_, 0.05% tween-80, cOmplete™ EDTA-free Protease Inhibitor. The cells were lysed by a double pass through a Stansted pressure cell homogeniser (270 MPa). Cell debris was pelleted and discarded by centrifugation at 10.000 rcf, 20 min at 4°C. The crude membranes were extracted by ultra-centrifugation (200.000 rcf, 1 hour, 4°C) and resuspended in 20 mM Tris-HCl, pH 7.4, 0.05% Tween-80, 10% glycerol to a total protein concentration of 10 mg/mL. The proteins were solubilized by the addition of 0.5% LMNG with gentle mixing (1h, 4°C). Insoluble material was pelleted by ultra-centrifugation (200.000 rcf, 30 min, 4°C) and discarded. The soluble fraction was incubated overnight at 4°C with Pierce™ Anti-DYKDDDDK Affinity Resin (Thermo Fisher) and purified according to the manufacturer’s instructions. Briefly, the flow through was removed by centrifugation at 1000 rcf, followed by four washing steps with 10 CV of 50 mM Tris-HCl (pH 7.4), 150 mM NaCl, 0.005% LMNG. The protein was eluted by the addition of 50 mM Tris-HCl (pH 7.4), 150 mM NaCl, 0.02% DDM, 1 mg/mL 3X-FLAG peptide (Genscript). Final purification was achieved by size exclusion chromatography (SEC) on a superdex 200 increase 10/200 column at 0.5 mL/min (50 mM Tris-HCl (pH 7.4), 150 mM NaCl, 0.005% LMNG). SEC peak fractions were evaluated on SDS-gel and pure *M. tuberculosis* cytochrome bd was pooled, concentrated, and stored at -80°C until further use.

### Expression and purification of NDH-2

The gene for *Caldalkalibacillus thermarum* NDH-2 with an N-terminal hexahistidine tag was ordered from GeneArt and cloned in the pET28 vector between NcoI and XhoI, giving rise to the construct pET28-NDH-2_NtermHis. pET28-NDH-2_NtermHis was transformed into C41 (DE3) cells and plated on LB kanamycin to select positive transformants.

Expression and purification were performed based on the procedure from Heikal et al. with slight modifications ^55^. Briefly, a streak of transformants was inoculated and grown overnight (250 RPM, 37°C). The overnight culture was diluted to ∼OD 0.1 in LB kanamycin and grown to ∼OD 0.5 before induction with 0.25 mM IPTG. Expression was carried out for 4 hours before cells were harvested (6371 rcf, 20 min, 4°C). The cells were resuspended in a 5-fold volume of 50 mM Tris-HCl, 5 mM MgCl_2_, pH 8.0, and lysed by a single pass through a Stansted pressure cell homogeniser (270 MPa). Unbroken cells were pelleted by centrifugation (10.000 rcf, 20 min, 4°C) and discarded. Crude membrane fractions were pelleted by ultra-centrifugation (200.000 rcf, 1h, 4°C) and resuspended at a 10 mg/mL total protein concentration in Tris-HCl, 150 mM NaCl, 20 mM Imidazole. Membrane proteins were extracted by treatment with 1% DDM for 1 hour at 4°C with gentle mixing. The membranes were removed by ultra-centrifugation (200.000 rcf, 30min, 4°C) followed by application of the soluble fraction to a HiTrap Nickel NTA column (Cytiva). The unbound proteins were washed from the column with washing buffer (50 mM Tris-HCl pH 8.0, 150 mM NaCl, 20 mM Imidazole, 0.02% DDM) followed by elution using stepwise addition of elution buffer (50 mM Tris-HCl, 150 mM NaCl, 500 mM Imidazole, 0.02% DDM). NDH-2 eluted at approximately 30% elution buffer, as confirmed by SDS-page gel and western blot. Final purification was achieved by size exclusion chromatography on a superdex increase 200 10/300 column (Cytiva) at 0.5 mL/min (50 mM Tris-HCl, 500 mM NaCl, 5% glycerol, 0.02% DDM). Pure fractions were pooled, concentrated, and stored at -80°C until further use.

### Cyt *bd* reconstitution in proteoliposomes

Lipids were purchased from Avanti Polar Lipids and used as received. A lipid mixture of POPE:POPG:CA (60:30:10 for *Ecbd* proteoliposomes, 30:60:10 for *Mtbd* proteoliposomes), enriched with the desired concentrations of ubiquinone-10 (UQ-10) (Sigma) or menaquinone-9 (MK-9) (Caymen chemical), was dried under a stream of nitrogen. Final traces of CHCl_3_ were removed overnight under vacuum. The lipid film was rehydrated and resuspended to a final concentration of 10 mg/mL in 20 mM MOPS, 30 mM Na_2_SO_4_, 100 mM KCl, pH 7,4, by vortexing. Cyt *bd* reconstitution was performed as described ^56^. Briefly, cyt *bd* was added to the liposome solution at 1 w/w% protein/lipids and mixed for 30 min by inversion at RT. Insoluble materials was removed by centrifugation in an Eppendorf tabletop centrifuge (14100 rcf, 5 min). The proteoliposome cyt *bd* concentration was determined by re-dissolving a sample in 2% octyl-β-glucoside, followed by quantification of the soret band with the corresponding extinction coefficient (*Ecbd*: ε_417_ 230 mM^-1^ cm^-1 57^, *Mtbd*: ε_414_ 279 mM^-1^ cm^-1^). The latter extinction coefficient of the *Mtbd* soret band (414 nm) was determined from UV vis absorbance in relation to protein concentrations determined by BCA assay from three different protein preparations. The extinction coefficient (ε_414nm_ = 279 ± 13 mM^-1^ cm^-1^) was comparable to other bd-type oxidases ^58^.

### Enzyme kinetics with water soluble quinone analogs

Oxygen consumption of LMNG solubilized cyt *bd* with quinone analogs was measured on an oxygraph (Hansatech Ltd.) system at 20°C. The quinone analogs, UQ-1 (Sigma), deoxylapachol (DMK-1, MedChem Express), or MK-1 (Santa Cruz Biotech) were added to the reaction chamber at the desired concentration in 50 mM MOPS, 150 mM NaCl, pH 7.0. Quinone-mediated autooxidation was determined by enzymatic reduction of the quinones by *C. thermarum* NDH-2 (30 nM) after the addition of 1 mM NADH. Oxygen consumption was initiated by the addition of cyt *bd* (4 nM for *Ecbd*, 6.5 nM for *Mtbd*). The enzyme activity was measured by subtraction of the quinone autooxidation rate from the initial slope after cyt *bd* addition. The kinetics curves were fit and where possible, enzymatic parameters were determined using GraphPad Prism using either Michaelis Menten (eq.1) or substrate inhibition kinetics (eq.2).

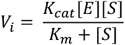

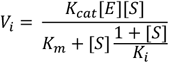

### Cyt *bd* oxygen consumption in proteoliposomes

Oxygen consumption was measured on an oxygraph system at 20°C. Prior to the measurement, cytochrome bd proteoliposomes were diluted to the desired concentration (20 nM cyt *bd*) in 50 mM MOPS, 150 mM NaCl, pH 7.0. The proteoliposomes were incubated for 30 min at room temperature with 100 nM NDH-2 to complete the proteoliposomal system. Oxygen consumption was initiated by the addition of 1 mM NADH. The oxygen consumption rate was quantified using the initial slope after NADH addition.

### Cyt *bd* oxygen consumption after treatment with reductants

The effect of *Mtbd* Q-loop disulfide bond reduction was studied by 30 minutes of preincubation with chemical reductants (10 mM TCEP, 10 mM DTT, 100 mM 2-ME). The samples were diluted to the concentrations used previously in the LMNG solubilized or proteoliposomal measurements. The buffer was supplemented with the respective chemical reductant to maintain the reductive environment during the measurement (1 mM TCEP, 10 mM DTT, 100 mM 2-ME). The measurements were performed as mentioned before.

## Acknowledgements

We would like to thank Dr. Dirk Bald (VU Amsterdam, The Netherlands) for providing the MB43 cells and the pET17b_CydABX_Linkerstreptag construct for *Ecbd* expression. Furthermore, we would like to thank Prof. Dr. Gregory Cook (University of Otago, New Zealand) for providing the MC^2^ 155 ΔCydAB strain and pLHCyd plasmid for *Mtbd* expression.

## Author contributions

Conceptualization: T.V. and L.J., Experiments and analysis: T.V., NDH-2 purification: T.V. and C.W., TV and LJ wrote the manuscript.

## Abbreviations

Cyt *bd*: Cytochrome *bd*
DMK-1: demethylmenaquinone-1
DTT: Dithiothreitol
*Ecbd*: *E. coli* cytochrome bd-I
LMNG: Lauryl Maltose Neopentyl Glycol
MK: Menaquinone
*Mtbd*: *M. tuberculosis* cytochrome bd
NDH-2: NADH dehydrogenase type-2
TCEP: Tris(2-carboxyethyl)phosphine hydrochloride
UQ: Ubiquinone-10
2-ME: 2-Mercaptoethanol

## References

(1) Belevich, I.; Borisov, V. B; Bloch, D. A.; Konstantinov, A. A.; Verkhovsky, M. I. Cytochrome Bd from Azotobacter Vinelandii: Evidence for High-Affinity Oxygen Binding. Biochemistry 2007, 46 (39), 11177–11184. 10.1021/bi700862u.

(2) D’mello’, R.; Hill’t, S.; Poolel, R. K. The Cytochrome Bd Quinol Oxidase in Escherichia Coli Has an Extremely High Oxygen Affinity and Two Oxygen-Binding Haems: Implications for Regulation of Activity in Vivo by Oxygen Inhibition. Microbiology (N Y) 1996, 142, 755–763. 10.1099/00221287-142-4-755.

(3) Borisov, V. B.; Siletsky, S. A; Paiardini, A.; Hoogewijs, D.; Forte, E.; Giuffrè, A.; Poole, R. K. Bacterial Oxidases of the Cytochrome Bd Family: Redox Enzymes of Unique Structure, Function, and Utility As Drug Targets. Antioxid Redox Signal 2021, 34 (16), 1280–1318. 10.1089/ars.2020.8039.

(4) Lee, B. S.; Hards, K; Engelhart, C. A.; Hasenoehrl, E. J.; Kalia, N. P.; Mackenzie, J. S.; Sviriaeva, E.; Chong, S. M. S.; Manimekalai, M. S. S.; Koh, V. H.; Chan, J.; Xu, J.; Alonso, S.; Miller, M. J.; Steyn, A. J. C.; Grüber, G.; Schnappinger, D.; Berney, M.; Cook, G. M.; Moraski, G. C.; Pethe, K. Dual Inhibition of the Terminal Oxidases Eradicates Antibiotic-tolerant Mycobacterium Tuberculosis. EMBO Mol Med 2021, 13 (1), 1–16. 10.15252/emmm.202013207.

(5) Kalia, N. P.; Hasenoehrl, E. J; Rahman, N. B. A.; Koh, V. H.; Ang, M. L. T.; Sajorda, D. R.; Hards, K.; Grüber, G.; Alonso, S.; Cook, G. M.; Berney, M.; Pethe, K. Exploiting the Synthetic Lethality between Terminal Respiratory Oxidases to Kill Mycobacterium Tuberculosis and Clear Host Infection. Proc Natl Acad Sci U S A 2017, 114 (28), 7426–7431. 10.1073/pnas.1706139114.

(6) Giuffrè, A.; Borisov, V. B.; Arese, M.; Sarti, P.; Forte, E. Cytochrome Bd Oxidase and Bacterial Tolerance to Oxidative and Nitrosative Stress. Biochim Biophys Acta Bioenerg 2014, 1837 (7), 1178–1187. 10.1016/j.bbabio.2014.01.016.

(7) Forte, E.; Borisov, V. B; Falabella, M.; Colaço, H. G.; Tinajero-Trejo, M.; Poole, R. K.; Vicente, J. B.; Sarti, P.; Giuffre, A. The Terminal Oxidase Cytochrome Bd Promotes Sulfide-Resistant Bacterial Respiration and Growth. Sci Rep 2016, 6. 10.1038/srep23788.

(8) Lu, P.; Heineke, M. H; Koul, A.; Andries, K.; Cook, G. M.; Lill, H.; Van Spanning, R.; Bald, D. The Cytochrome Bd-Type Quinol Oxidase Is Important for Survival of Mycobacterium Smegmatis under Peroxide and Antibiotic-Induced Stress. Sci Rep 2015, 5. 10.1038/srep10333.

(9) Beebout, C. J.; Sominsky, L. A; Eberly, A. R.; Van Horn, G. T.; Hadjifrangiskou, M. Cytochrome Bd Promotes Escherichia Coli Biofilm Antibiotic Tolerance by Regulating Accumulation of Noxious Chemicals. biofilms and Microbiomes 2021, 7 (1). 10.1038/s41522-021-00210-x.

(10) Seregina, T. A.; Lobanov, K. V; Shakulov, R. S.; Mironov, A. S. Inactivation of Terminal Oxidase Bd-I Leads to Supersensitivity of E. Coli to Quinolone and Beta-Lactam Antibiotics. Mol Biol 2022, 56 (4), 572–579. 10.1134/S0026893322040100.

(11) Xia, X.; Wu, S; Li, L.; Xu, B.; Wang, G. The Cytochrome Bd Complex Is Essential for Chromate and Sulfide Resistance and Is Regulated by a GbsR-Type Regulator, CydE, in Alishewanella Sp. WH16-1. Front Microbiol 2018, 9 (1849). 10.3389/fmicb.2018.01849.

(12) Safarian, S.; Opel-Reading, H. K; Wu, D.; Mehdipour, A. R.; Hards, K.; Harold, L. K.; Radloff, M.; Stewart, I.; Welsch, S.; Hummer, G.; Cook, G. M.; Krause, K. L.; Michel, H. The Cryo-EM Structure of the Bd Oxidase from M. Tuberculosis Reveals a Unique Structural Framework and Enables Rational Drug Design to Combat TB. Nat Commun 2021, 12 (1). 10.1038/s41467-021-25537-z.

(13) Grund, T. N.; Kabashima, Y; Kusumoto, T.; Wu, D.; Welsch, S.; Sakamoto, J.; Michel, H.; Safarian, S. The CryoEM Structure of Cytochrome Bd from C. Glutamicum Provides Novel Insights into Structural Properties of Actinobacterial Terminal Oxidases. Front Chem 2023, 10. 10.3389/fchem.2022.1085463.

(14) Safarian, S.; Hahn, A; Mills, D. J.; Radloff, M.; Eisinger, M. L.; Nikolaev, A.; Meier-Credo, J.; Melin, F.; Miyoshi, H.; Gennis, R. B.; Sakamoto, J.; Langer, J. D.; Hellwig, P.; Kühlbrandt, W.; Michel, H. Active Site Rearrangement and Structural Divergence in Prokaryotic Respiratory Oxidases. Science 2019, 366 (6461), 100–104. 10.1126/science.aay0967.

(15) Mogi, T.; Akimoto, S; Endou, S.; Watanabe-Nakayama, T.; Mizuochi-Asai, E.; Miyoshi, H. Probing the Ubiquinol-Binding Site in Cytochrome Bd by Site-Directed Mutagenesis. Biochemistry 2006, 45 (25), 7924–7930. 10.1021/bi060192w.

(16) Grund, T. N.; Radloff, M; Wu, D.; Goojani, H. G.; Witte, L. F.; Jösting, W.; Buschmann, S.; Müller, H.; Elamri, I.; Welsch, S.; Schwalbe, H.; Michel, H.; Bald, D.; Safarian, S. Mechanistic and Structural Diversity between Cytochrome Bd Isoforms of Escherichia Coli. Proc Natl Acad Sci U S A 2021, 118 (50). 10.1073/pnas.2114013118.

(17) Harikishore, A.; Mathiyazakan, V; Pethe, K.; Grüber, G. Novel Targets and Inhibitors of the Mycobacterium Tuberculosis Cytochrome Bd Oxidase to Foster Anti-Tuberculosis Drug Discovery. Expert Opin Drug Discov 2023, 18 (8), 917–927. 10.1080/17460441.2023.2224553.

(18) Cremers, C. M.; Jakob, U. Oxidant Sensing by Reversible Disulfide Bond Formation. Journal of Biological Chemistry 2013, 288 (37), 26489–26496. 10.1074/jbc.R113.462929.

(19) Oliveira, A. R.; Mota, C; Vilela-Alves, G.; Manuel, R. R.; Pedrosa, N.; Fourmond, V.; Klymanska, K.; Léger, C.; Guigliarelli, B.; Romão, M. J.; Cardoso Pereira, I. A. An Allosteric Redox Switch Involved in Oxygen Protection in a CO2 Reductase. Nat Chem Biol 2023. 10.1038/s41589-023-01484-2.

(20) Unden, G.; Bongaerts, J. Alternative Respiratory Pathways of Escherichia Coli: Energetics and Transcriptional Regulation in Response to Electron Acceptors. Biochim Biophys Acta 1997, 1320, 217–234. 10.1016/s0005-2728(97)00034-0.

(21) Kaurola, P.; Sharma, V; Vonk, A.; Vattulainen, I.; Róg, T. Distribution and Dynamics of Quinones in the Lipid Bilayer Mimicking the Inner Membrane of Mitochondria. Biochim Biophys Acta Biomembr 2016, 1858 (9), 2116–2122. 10.1016/j.bbamem.2016.06.016.

(22) Collins, M. D.; Jones, D. Distribution of Isoprenoid Quinone Structural Types in Bacteria and Their Taxonomic Implications. Microbiol Rev 1981, 45 (2), 316–354. 10.1128/mr.45.2.316-354.1981.

(23) Sharma, P.; Teixeira De Mattos, M. J; Hellingwerf, K. J.; Bekker, M. On the Function of the Various Quinone Species in Escherichia Coli. FEBS Journal 2012, 279 (18), 3364–3373. 10.1111/j.1742-4658.2012.08608.x.

(24) van Beilen, J. W. A.; Hellingwerf, K. J. All Three Endogenous Quinone Species of Escherichia Coli Are Involved in Controlling the Activity of the Aerobic/Anaerobic Response Regulator ArcA. Front Microbiol 2016, 7 (1339). 10.3389/fmicb.2016.01339.

(25) Borisov, V. B.; Gennis, R. B; Hemp, J.; Verkhovsky, M. I. The Cytochrome Bd Respiratory Oxygen Reductases. Biochim Biophys Acta Bioenerg 2011, 1807 (11), 1398–1413. 10.1016/j.bbabio.2011.06.016.

(26) Lamprecht, D. A.; Finin, P. M; Rahman, M. A.; Cumming, B. M.; Russell, S. L.; Jonnala, S. R.; Adamson, J. H.; Steyn, A. J. C. Turning the Respiratory Flexibility of Mycobacterium Tuberculosis against Itself. Nat Commun 2016, 7. 10.1038/ncomms12393.

(27) Kusumoto, K.; Sakiyama, M; Sakamoto, J.; Noguchi, S.; Sone, N. Menaquinol Oxidase Activity and Primary Structure of Cytochrome Bd from the Amino-Acid Fermenting Bacterium Corynebacterium Glutamicum. Arch Microbiol 2000, 173 (5–6), 390–397. 10.1007/s002030000161.

(28) Lorence, R. M.; Miller, M. J; Borochov, A.; Faiman-Weinberg, R.; Gennis, R. B. Effects of PH and Detergent on the Kinetic and Electrochemical Properties of the Purified Cytochrome d Terminal Oxidase Complex of Escherichia Coli. Biochim Biophys Acta 1984, 790, 148. 10.1016/0167-4838(84)90218-8.

(29) Upadhyay, A.; Fontes, F. L; Gonzalez-Juarrero, M.; McNeil, M. R.; Crans, D. C.; Jackson, M.; Crick, D. C. Partial Saturation of Menaquinone in Mycobacterium Tuberculosis: Function and Essentiality of a Novel Reductase, MenJ. ACS Cent Sci 2015, 1 (6), 292–302. 10.1021/acscentsci.5b00212.

(30) Yanofsky, D. J.; Di Trani, J. M; Krol, S.; Abdelaziz, R.; Bueler, S. A.; Imming, P.; Brzezinski, P.; Rubinstein, J. L. Structure of Mycobacterial CIII2CIV2 Respiratory Supercomplex Bound to the Tuberculosis Drug Candidate Telacebec (Q203). Elife 2021, 10 (e71959). 10.7554/eLife.71959.

(31) Matsumoto, Y.; Muneyuki, E; Fujita, D.; Sakamoto, K.; Miyoshi, H.; Yoshida, M.; Moggi, T. Kinetic Mechanism of Quinol Oxidation by Cytochrome Bd Studied with Ubiquinone-2 Analogs. J Biochem 2006, 139 (4), 779–788. 10.1093/jb/mvj087.

(32) Sakamoto, K.; Miyoshi, H; Takegami, K.; Mogi, T.; Anraku, Y.; Iwamura, H. Probing Substrate Binding Site of the Escherichia Coli Quinol Oxidases Using Synthetic Ubiquinol Analogues. Journal of Biological Chemistry 1996, 271 (47), 29897–29902. 10.1074/jbc.271.47.29897.

(33) Velappan, A. B.; Datta, D; Ma, R.; Rana, S.; Ghosh, K. S.; Hari, N.; Franzblau, S. G.; Debnath, J. 2-Aryl Benzazole Derived New Class of Anti-Tubercular Compounds: Endowed to Eradicate Mycobacterium Tuberculosis in Replicating and Non-Replicating Forms. Bioorg Chem 2020, 103. 10.1016/j.bioorg.2020.104170.

(34) Goddard, T. D.; Huang, C. C; Meng, E. C.; Pettersen, E. F.; Couch, G. S.; Morris, J. H.; Ferrin, T. E. UCSF ChimeraX: Meeting Modern Challenges in Visualization and Analysis. Protein Science 2018, 27 (1), 14–25. 10.1002/pro.3235.

(35) Kita, K.; Konishi, K; Anraku, Y. Terminal Oxidases of Escherichia Coli Aerobic Respiratory Chain. II. Purification and Properties of Cytochrome B558-d Complex from Cells Grown with Limited Oxygen and Evidence of Branched Electron-Carrying Systems. Journal of Biological Chemistry 1984, 259 (5), 3375–3381. 10.1016/s0021-9258(17)43305-9.

(36) Asseri, A. H.; Godoy-Hernandez, A; Goojani, H. G.; Lill, H.; Sakamoto, J.; McMillan, D. G. G.; Bald, D. Cardiolipin Enhances the Enzymatic Activity of Cytochrome Bd and Cytochrome Bo 3 Solubilized in Dodecyl-Maltoside. Sci Rep 2021, 11 (1). 10.1038/s41598-021-87354-0.

(37) Miller, M. J.; Gennis, R. B. The Purification and Characterization of the Cytochrome d Terminal Oxidase Complex of the Escherichia Coli Aerobic Respiratory Chain*. Journal of Biological Chemistry 1983, 258 (15), 9159–9165. 10;258(15):9159-65.

(38) Shestopalov, A. I.; Bogachev, A. V; Murtazina, R. A.; Viryasov, M. B.; Skulachev, V. P. Aeration-Dependent Changes in Composition of the Quinone Pool in Escherichia Coli. FEBS Lett 1997, 404 (2–3), 272–274. 10.1016/S0014-5793(97)00143-9.

(39) Jeffreys, L. N.; Ardrey, A; Hafiz, T. A.; Dyer, L. A.; Warman, A. J.; Mosallam, N.; Nixon, G. L.; Fisher, N. E.; Hong, W. D.; Leung, S. C.; Aljayyoussi, G.; Bibby, J.; Almeida, D. V.; Converse, P. J.; Fotouhi, N.; Berry, N. G.; Nuermberger, E. L.; Upton, A. M.; O’Neill, P. M.; Ward, S. A.; Biagini, G. A. Identification of 2-Aryl-Quinolone Inhibitors of Cytochrome Bd and Chemical Validation of Combination Strategies for Respiratory Inhibitors against Mycobacterium Tuberculosis. ACS Infect Dis 2023, 9 (2), 221–238. 10.1021/acsinfecdis.2c00283.

(40) Li, J.; Han, L; Vallese, F.; Ding, Z.; Choi, S. K.; Hong, S.; Luo, Y.; Liu, B.; Chan, C. K.; Tajkhorshid, E.; Zhu, J.; Clarke, O.; Zhang, K.; Gennis, R. Cryo-EM Structures of Escherichia Coli Cytochrome Bo3 Reveal Bound Phospholipids and Ubiquinone-8 in a Dynamic Substrate Binding Site. Proc Natl Acad Sci U S A 2021, 118 (34). 10.1073/pnas.2106750118.

(41) Nakatani, Y.; Shimaki, Y; Dutta, D.; Muench, S. P.; Ireton, K.; Cook, G. M.; Jeuken, L. J. C. Unprecedented Properties of Phenothiazines Unraveled by a NDH-2 Bioelectrochemical Assay Platform. J Am Chem Soc 2020, 142 (3), 1311–1320. 10.1021/jacs.9b10254.

(42) Nikolaev, A.; Safarian, S; Thesseling, A.; Wohlwend, D.; Friedrich, T.; Michel, H.; Kusumoto, T.; Sakamoto, J.; Melin, F.; Hellwig, P. Electrocatalytic Evidence of the Diversity of the Oxygen Reaction in the Bacterial Bd Oxidase from Different Organisms. Biochim Biophys Acta Bioenerg 2021, 1862 (8). 10.1016/j.bbabio.2021.148436.

(43) Kao, W. C.; Kleinschroth, T; Nitschke, W.; Baymann, F.; Neehaul, Y.; Hellwig, P.; Richers, S.; Vonck, J.; Bott, M.; Hunte, C. The Obligate Respiratory Supercomplex from Actinobacteria. Biochim Biophys Acta Bioenerg 2016, 1857 (10), 1705–1714. 10.1016/j.bbabio.2016.07.009.

(44) Sukheja, P.; Kumar, P; Mittal, N.; Li, S. G.; Singleton, E.; Russo, R.; Perryman, A. L.; Shrestha, R.; Awasthi, D.; Husain, S.; Soteropoulos, P.; Brukh, R.; Connell, N.; Freundlich, J. S.; Alland, D. A Novel Small-Molecule Inhibitor of the Mycobacterium Tuberculosis Demethylmenaquinone Methyltransferase Meng Is Bactericidal to Both Growing and Nutritionally Deprived Persister Cells. mBio 2017, 8 (1). 10.1128/mBio.02022-16.

(45) Wang, W.; Gao, Y; Tang, Y.; Zhou, X.; Lai, Y.; Zhou, S.; Zhang, Y.; Yang, X.; Liu, F.; Guddat, L. W.; Wang, Q.; Rao, Z.; Gong, H. Cryo-EM Structure of Mycobacterial Cytochrome Bd Reveals Two Oxygen Access Channels. Nat Commun 2021, 12 (1). 10.1038/s41467-021-24924-w.

(46) Voskuil, M. I.; Bartek, I. L; Visconti, K.; Schoolnik, G. K. The Response of Mycobacterium Tuberculosis to Reactive Oxygen and Nitrogen Species. Front Microbiol 2011, 2 (105). 10.3389/fmicb.2011.00105.

(47) Small, J. L.; Park, S. W; Kana, B. D.; Ioerger, T. R.; Sacchettini, J. C.; Ehrt, S. Perturbation of Cytochrome c Maturation Reveals Adaptability of the Respiratory Chain in Mycobacterium Tuberculosis. mBio 2013, 4 (5), 475–488. 10.1128/MBIO.00475-13/ASSET/DAECB6C2-C85B-4593-A9C4-7C49664B4B16/ASSETS/GRAPHIC/MBO0051316240004.JPEG.

(48) Al-Attar, S.; Yu, Y; Pinkse, M.; Hoeser, J.; Friedrich, T.; Bald, D.; De Vries, S. Cytochrome Bd Displays Significant Quinol Peroxidase Activity. Sci Rep 2016, 6. 10.1038/srep27631.

(49) Borisov, V. B.; Forte, E; Siletsky, S. A.; Sarti, P.; Giuffrè, A. Cytochrome Bd from Escherichia Coli Catalyzes Peroxynitrite Decomposition. Biochim Biophys Acta Bioenerg 2015, 1847 (2), 182–188. 10.1016/j.bbabio.2014.10.006.

(50) Goojani, H. G.; Konings, J; Hakvoort, H.; Hong, S.; Gennis, R. B.; Sakamoto, J.; Lill, H.; Bald, D. The Carboxy-Terminal Insert in the Q-Loop Is Needed for Functionality of Escherichia Coli Cytochrome Bd-I. Biochim Biophys Acta Bioenerg 2020, 1861 (5–6). 10.1016/j.bbabio.2020.148175.

(51) Bekker, M.; De Vries, S; Ter Beek, A.; Hellingwerf, K. J.; Teixeira De Mattos, M.J. Respiration of Escherichia Coli Can Be Fully Uncoupled via the Nonelectrogenic Terminal Cytochrome Bd-II Oxidase. J Bacteriol 2009, 191 (17), 5510–5517. 10.1128/JB.00562-09.

(52) Arai, H.; Kawakami, T; Osamura, T.; Hirai, T.; Sakai, Y.; Ishii, M. Enzymatic Characterization and in Vivo Function of Five Terminal Oxidases in Pseudomonas Aeruginosa. J Bacteriol 2014, 196 (24), 4206–4215. 10.1128/JB.02176-14.

(53) Safarian, S.; Opel-Reading, H. K; Wu, D.; Mehdipour, A. R.; Hards, K.; Harold, L. K.; Radloff, M.; Stewart, I.; Welsch, S.; Hummer, G.; Cook, G. M.; Krause, K. L.; Michel, H. The Cryo-EM Structure of the Bd Oxidase from M. Tuberculosis Reveals a Unique Structural Framework and Enables Rational Drug Design to Combat TB. Nat Commun 2021, 12 (1). 10.1038/s41467-021-25537-z.

(54) Lu, X.; Williams, Z; Hards, K.; Tang, J.; Cheung, C. Y.; Aung, H. L.; Wang, B.; Liu, Z.; Hu, X.; Lenaerts, A.; Woolhiser, L.; Hastings, C.; Zhang, X.; Wang, Z.; Rhee, K.; Ding, K.; Zhang, T.; Cook, G. M. Pyrazolo[1,5-a]Pyridine Inhibitor of the Respiratory Cytochrome Bcc Complex for the Treatment of Drug-Resistant Tuberculosis. ACS Infect Dis 2019, 5 (2), 239–249. 10.1021/acsinfecdis.8b00225.

(55) Heikal, A.; Nakatani, Y; Dunn, E.; Weimar, M. R.; Day, C. L.; Baker, E. N.; Lott, J. S.; Sazanov, L. A.; Cook, G. M. Structure of the Bacterial Type II NADH Dehydrogenase: A Monotopic Membrane Protein with an Essential Role in Energy Generation. Mol Microbiol 2014, 91 (5), 950–964. 10.1111/mmi.12507.

(56) Godoy-Hernandez, A.; Asseri, A. H; Purugganan, A. J.; Jiko, C.; de Ram, C.; Lill, H.; Pabst, M.; Mitsuoka, K.; Gerle, C.; Bald, D.; McMillan, D. G. G. Rapid and Highly Stable Membrane Reconstitution by LAiR Enables the Study of Physiological Integral Membrane Protein Functions. ACS Cent Sci 2023, 9 (3), 494–507. 10.1021/acscentsci.2c01170.

(57) Tsubaki, M.; Hori, H; Mogi, T.; Anraku, Y. Cyanide-Binding Site of Bd-Type Ubiquinol Oxidase from Escherichia Coli*. Journal of Biological Chemistry 1995, 270 (48), 28565–28569.

(58) Borisov, V. B.; Gennis, R. B; Hemp, J.; Verkhovsky, M. I. The Cytochrome Bd Respiratory Oxygen Reductases. Biochim Biophys Acta Bioenerg 2011, 1807 (11), 1398–1413. 10.1016/j.bbabio.2011.06.016.

