## Supplementary information for "Menaquinone-specific oxidation by *M. tuberculosis* cytochrome *bd* is redox regulated by the Q-loop disulfide bond"

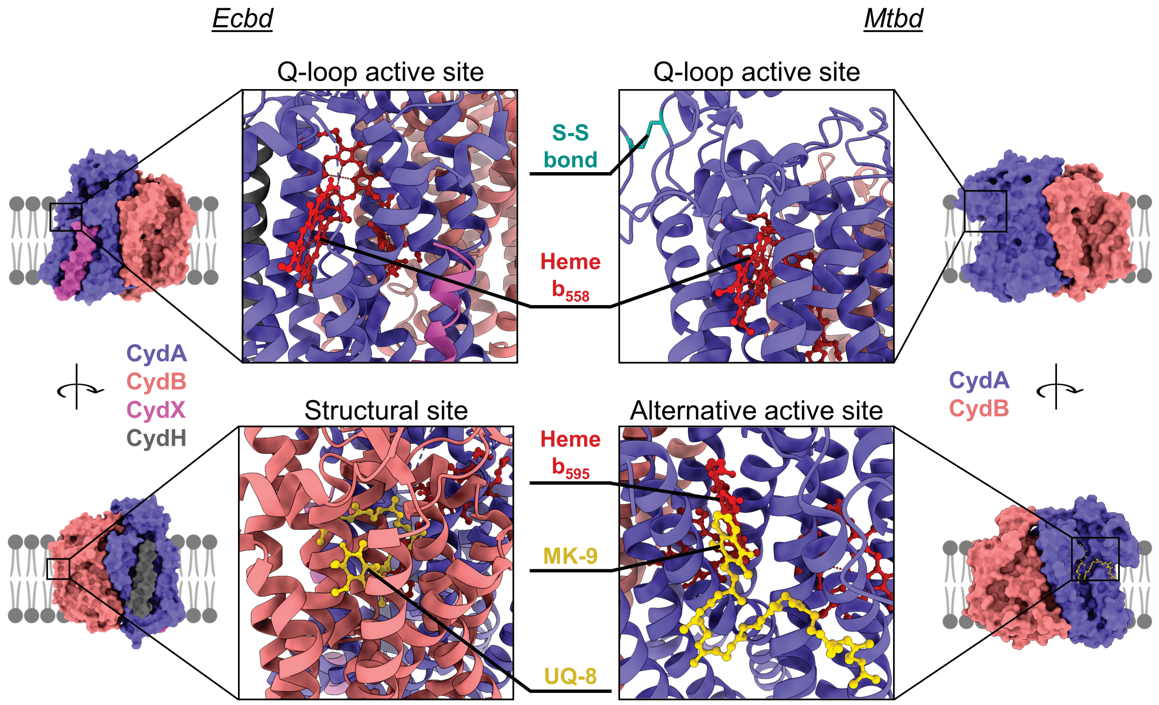


**SI figure 1: Cryo EM strucures of *Ecbd* (PDB: 6RKO) and *Mtbd* (PDB: 7NKZ) with active and structural sites indicated.**

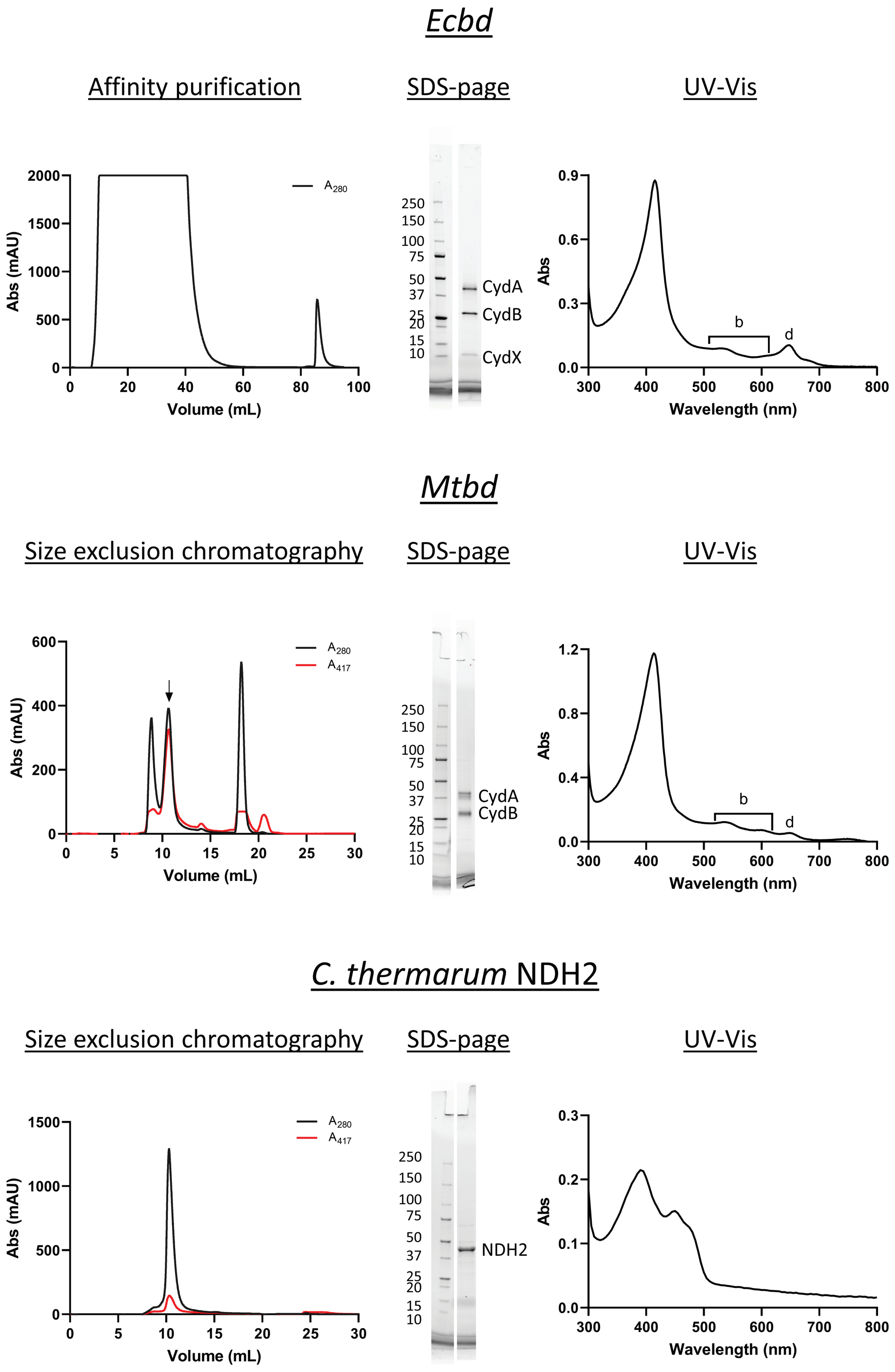


**SI figure 2.** Purification, SDS-Page, and UV-Vis spectra of the isolated *Ecbd*, *Mtbd* and *C. thermarum* NDH2.

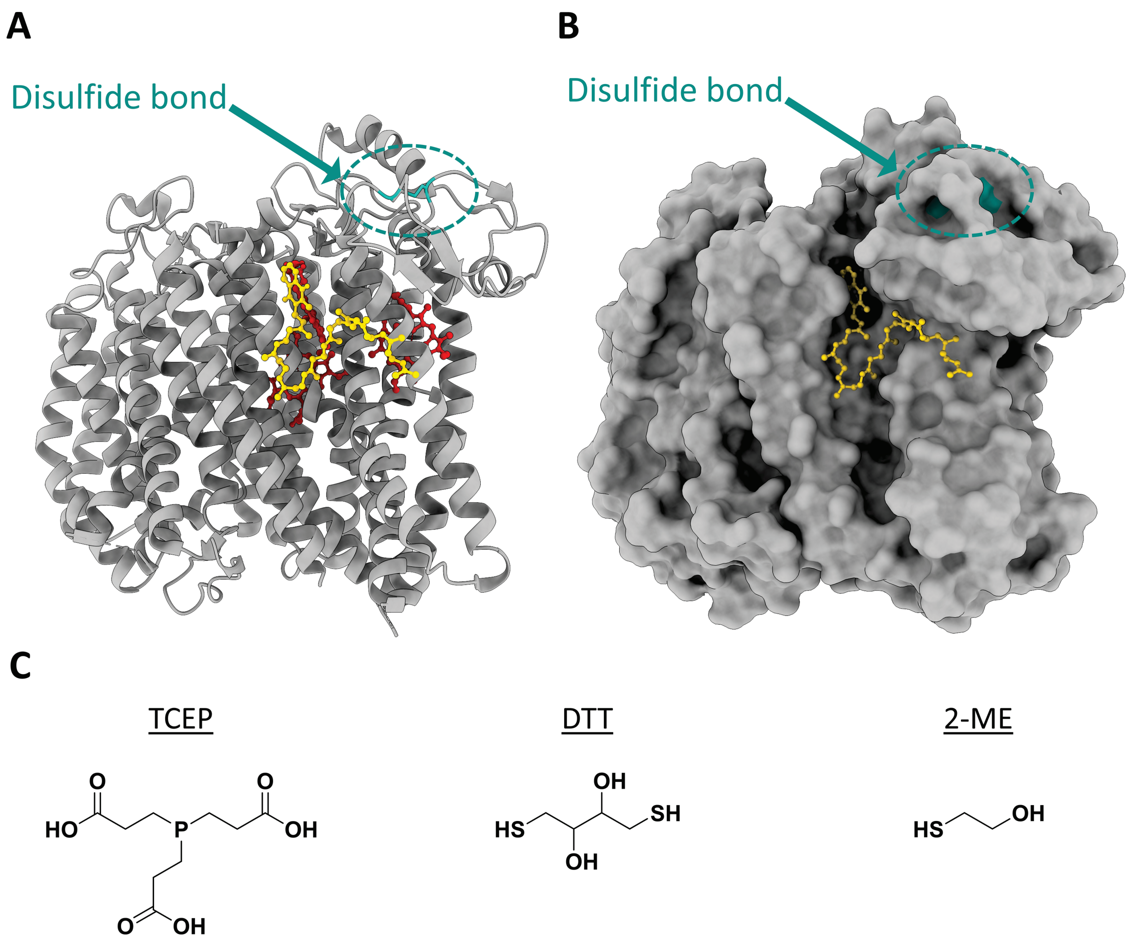


**SI figure 3**. Accessibility of the *Mtbd* disulfide bond. Cartoon (**A**) and surface (**B**) representation of *Mtbd* with hemes (Red), MK-9 (Blue) and disulfide bond (Purple). (**C**) chemical structure of the reductants used in this study.

**Calculation of *E. coli* membrane quinone content**

| Variables | Value | Ref |
|---|---|---|
| UQ content *E. coli* | $1.5 \times 10^{3}$ nmol/g dry cell weight | 1 |
| | $3.6 \times 10^{2}$ nmol/g dry cell weight | 2 |
| *E. coli* dry weight | $3.0 \times 10^{-13}$ gram per cell | 3 |
| Number of lipids per *E. coli* cell | $2.2 \times 10^{7}$ lipids | 3 |
| Avogadro's number | $6.022 \times 10^{23}$ | |

<u>Calculation</u>

Number of lipid per cell: $2.2 \times 10^{7}$ / $6.02 \times 10^{23}$ = $3.6 \times 10^{-17}$ mol/cell

Number of lipids in 1 gram cells: $3.6 \times 10^{-17}$ mol/cell × ($1/3.0 \times 10^{-13}$ g/cell) = $1.2 \times 10^{5}$ nmol/g

% UQ in membrane[1]: = $1.5 \times 10^{3}$ nmol/g / $1.2 \times 10^{5}$ nmol/g × 100% = 1.23%

% UQ in membrane[2]: = $3.6 \times 10^{2}$ nmol/g / $1.2 \times 10^{5}$ nmol/g × 100% = 0.30%

Assuming a surface area of POPC to be 65 $Å^2$ (MW = 760 g/mol) and a lipid bilayer thickness of 4 nm, 1% (w/w) UQ-10 (MW = 865 g/mol) is equivalent to 11 mM

Volume of 1 mol POPC bilayer:

$(6.022 \times 10^{23} \times 0.65\ nm^2 \times 4\ nm) / 2 = 7.8 \times 10^{23}\ nm^3 = 0.8\ L$

1% (w/w) UQ-10 of 1 mol POPC is = 0.01 × 760 g = 7.6 g or (7.6 g / 865 g/mol) = 8.8 mmol

8.8 mmol / 0.8 L = 11 mM

### Supplementary references


(1) Sharma, P.; Teixeira De Mattos, M. J.; Hellingwerf, K. J.; Bekker, M. On the Function of the Various Quinone Species in Escherichia Coli. In *FEBS Journal*; 2012; Vol. 279, pp 3364–3373. https://doi.org/10.1111/j.1742-4658.2012.08608.x.

(2) Unden, G.; Bongaerts, J. Alternative Respiratory Pathways of Escherichia Coli: Energetics and Transcriptional Regulation in Response to Electron Acceptors. *Biochim Biophys Acta* **1997**, *1320*, 217–234. https://doi.org/10.1016/s0005-2728(97)00034-0.

(3) Sajed, T.; Marcu, A.; Ramirez, M.; Pon, A.; Guo, A. C.; Knox, C.; Wilson, M.; Grant, J. R.; Djoumbou, Y.; Wishart, D. S. ECMDB 2.0: A Richer Resource for Understanding the Biochemistry of E. Coli. *Nucleic Acids Res* **2016**, *44* (D1), D495–D501. https://doi.org/10.1093/nar/gkv1060.
